# jFuzzyMachine – An Open–source Fuzzy Logic–based Regulatory Inference Engine for High–throughput Biological Data

**DOI:** 10.1101/2020.10.06.315994

**Authors:** Paul Aiyetan

## Abstract

Elucidating mechanistic relationships between and among intracellular macromolecules is fundamental to understanding the molecular basis of normal and diseased processes. Here, we introduce `jFuzzyMachine` – a fuzzy logic-based regulatory network inference engine for high-throughput biological data. We describe its design and implementation. We demonstrate its functions on a sampled expression profile of the vorinostat-resistant HCT116 cell line. We compared `jFuzzyMachine`’s inferred regulatory network to that inferred by the `ARACNe` (an Algorithm for the Reconstruction of Gene Regulatory Networks) tool. Potentially more sensitive, `jFuzzyMachine` showed a slight increase in identified regulatory edges compared to `ARACNe`. A significant overlap was also observed in the identified edges between the two inference methods. Over 70 percent of edges identified by `ARACNe` were identified by `jFuzzyMachine`. Beyond identifying edges, `jFuzzyMachine` shows direction of interactions, including bidirectional interactions – specifying regulatory inputs and outputs of inferred relationships. `jFuzzyMachine` addresses an apparent lack of freely available community tool implementing a fuzzy logic regulatory network inference method – mitigating a limitation to applying and extending benefits of the fuzzy inference system to understanding biological data. `jFuzzyMachine`’s source codes and precompiled binaries are freely available at the Github repository locations:

https://github.com/paiyetan/jfuzzymachine and

https://github.com/paiyetan/jfuzzymachine/releases/tag/v1.7.21.

## 1 BACKGROUND

Elucidating mechanistic relationships among intracellular macromolecules is fundamental to understanding the molecular basis of normal and diseased processes. Traditional approaches to elucidating these involved low-throughput methods of obtaining reaction kinetics with attending resolution of associated partial or ordinary differential equations (ODEs) [6] [3][5,21,42]. However, such approaches being highly sensitive to quantitative errors and increased complexities when the inferential problem involves multiple parameters are subject to inaccurate analytical estimates, with respect to experimental data from high-throughput profiling [14, 37]. Partly to address these, the advent of high-throughput profiling approaches has been greeted with the development of reverse engineering and computational analyses methods that attempt to tease mechanistic relationships from attendant high-throughput data [6,11,45][13,35]. Amongst these, the classical fuzzy logic approach employs varying degrees of truth to describe relationships between interacting molecules. The fuzzy logic reasoning methods [48], having had significant successes in the design and implementation of high-end control systems [47, 49, 50], were first applied to biological data by Woolf and Wang [46]. In the subsequent years, methods have been proposed to address identified shortfalls associated with the original implementation and to improve the inferential capability of the fuzzy approach [**?, ?**, 4, 7, 12, 16, 24, 25, 31–34, 43, 44][38][36][9, 15]. In spite of advances in the theoretical basis and relative biological validity of the fuzzy approach, there exists the very apparent lack of analytical tools [1] that implement these methods available to the scientific and research community. The apparent lack of ready and freely available community tools limits applying the fuzzy inference approach to biological data. This also limits necessary comparisons and bench-marking of results obtained by the method against those obtained from comparable methods. To elucidate mechanistic relationships, and to make more readily available the Fuzzy logic inference approach, we developed the `jFuzzyMachine` as a freely available tool.

## 2 MATERIALS AND METHODS

### 2.1 Dataset

To demonstrate `jFuzzyMachine`’s functionalities, including comparing results with similar computational tool and to show how `jFuzzyMachine` may be used to infer regulatory network interactions among a population of genes, we downloaded and derived a sampled expression profile from the NCBI GEO Dataset GSE56788. The dataset is detailed under the BioProject accession PRJNA244587. This consists of a total of 45 assay samples from 15 biosamples, each ran in 3 independent biological replicates. RNA-seq expression profiles were acquired by next-generation sequencing of vorinostat-resistant HCT116 cells, following knockdown of potential vorinostat-resistant candidate genes. Expression profiles were compared to mock transfection (control). The authors of the study sought to understand the mechanisms by which these knockdowns contributed to vorinostat response. They employed the siRNA-mediated knockdown of each of some previously identified resistance candidate genes in the HCT116-VR cell line. RNA sequence expression data were downloaded from the NCBI Sequence Read Archive [19, 20], with accession number SRP041162.

#### 2.1.1 RNA Sequence Analyses – quality assessment, preprocessing and normalization

For quality assessment (QA) of curated fastq files, the `fastqcr, ngsReports and Rqc` R/bioconductor tools [17] [10, 28, 30], modeled after the FASTQC [2] tool philosophy were used. To quantify expression, sequence reads were aligned to the genome (NCBI GRCh38 build) using the TopHat2 [18] [40, 41] tool. Index files were downloaded from Illumina iGenomes archive. Accepted hits and annotation information in the BAM format [39] output files were assembled into an expression matrix of feature counts using the `featureCount` routine in the `Rsubread` package [22]. Feature counts were normalized using the DESeq2 package [23] tools-implemented regularized-log transformation to account for disparate total read counts in the different files and to allow for comparison across the different samples. For demonstration and evaluation purpose, the expression profile of 14 features were extracted. These include those for the genes PTHLH, KRT86, RUBCNL, CYS1, LINC00707, LINC00634, LINC00886, GCNT4, TAGLN3, ROBO4, MGAM, 2orf78, LOC105371789, and SERPINB7.

### 2.2 The fuzzy Logic Inference

Given an expression profile, the fuzzy logic approach involves three major steps:

- Fuzzification
- Rule evaluation, and
- Defuzzification

The fuzzification step derives qualitative values from the expression profile’s crisp values. It is often described as a mapping of non-fuzzy inputs to fuzzy linguistic terms [29]. Given qualitative values of HIGH, MEDIUM, or LOW, the fuzzification step takes a feature’s expression value and assigns it degrees to which it belongs to the respective class of HIGH, MEDIUM or LOW expression values. The rule evaluation step takes combinations of features and utilizes an inference engine rules of the form IF-THEN, including fuzzy set operations such as AND, OR, or NOT to evaluate input features in relation to outputs features. This has been described as attempting to make an expert judgment of collective linguistic terms; attempts to find a solution to an evaluation of the concurrent state of existence of linguistic description of states. The defuzzification step attempts to report a corresponding continuous numerical variable from a fuzzy state linguistic variable.

`jFuzzyMachine` implements the fuzzy inference steps described in [38][36][9, 15]. After an initial data transformation of log2 expression ratios by the `arctan` function and dividing values by 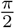, to project the ratios onto [*-*1, 1], the fuzzification step utilizes a membership function consisting of three fuzzy sets (low, medium, and high expression). Given three fuzzy sets (*y*_1_ = low, *y*_2_ = medium, *y*_3_ = high), its fuzzification of a gene expression value *x* results in the generation of a fuzzy set *y* = [*y*_1_, *y*_2_, *y*_3_] as follows:

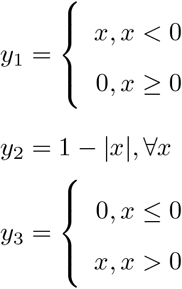

We specified our rule configuration (the specification of if-then relationships between variables in fuzzy space) in the form of a vector *r* = [*r*_1_, *r*_2_, *r*_3_]. We specified the state of an output node *z* = [*z*_1_, *z*_2_, *z*_3_] to be determined by the fuzzy state of an input feature *y* = [*y*_1_, *y*_2_, *y*_3_] and the rule describing the relationship between the input and the output, *r* = [*r*_1_, *r*_2_, *r*_3_] as follows:

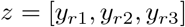

An inhibitory relationship, for example, specified as [3, 2, 1] implies, if input is low (*r*_1_), then output is high (3); if input is medium (*r*_2_), then output is medium (2), and if input is high (*r*_3_), then output in low (1). Classic fuzzy logic rule evaluation using the logical AND connective results in a combinatorial rule explosion i.e. an exponential increase in the number of rules to be evaluated and computational time, with additional inputs to be considered [8]. To address a combinatorial rule explosion situation, `jFuzzyMachine` implements the logical OR (union) rule configuration, an algebraic sum in fuzzy logic [48][27] as described in [37].

For Defuzzification, a conversion of fuzzy inferences or output node expression value predictions, in fuzzy space, back to regular or crisp values, we employed the simplified centroid method [27]. Given a predicted fuzzy values of an output node *y* = [*y*_1_, *y*_2_, *y*_3_], we defined defuzzified expression values 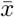 as:

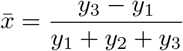

After defuzzification, we reverse transformed back to log2 expression values by multiplying derived values by 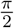 and applying the tangent function.

#### 2.2.1 Inferred regulatory model fit or error

`jFuzzyMachine` estimates the an error of the fit for *M* samples or perturbations of an output feature *x* = *{x*_1_, *x*_2_, …, *x*_*M*_ *}* as:

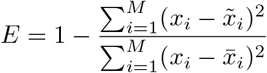

where 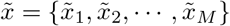 is the set of defuzzified numerical log expression ratios predicted for the output feature and 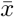 is the mean of the experimental values of *x* across the samples or perturbations observed. A perfect fit would result in a maximum *E* of 1.0. `jFuzzyMachine` uses the estimated error to rank possible models of interactions that predict on output node.

### 2.3 Design and Implementation

The `jFuzzyMachine` tool is implemented in the platform-independent Java programming language to facilitate an extensive community reach. It is modular in design to facilitate an easy decoupling of component parts. It consists of: 1) the Initiation Module, 2) the Main Module, and 3) the Utilities Module (see Figure 1). The ‘Initiation Module’ consists essentially of the program configuration and run parameter specification units. Depending on user-desired added-functionality beyond regulatory model inference, a user may choose to specify parameters that apply only to desired post-inference processing. The Main Module houses the application’s main functionality – the fuzzy logic based regulatory inference engine. The module implements the: fuzzification, rule evaluation, and defuzzification schemes [31, 46][29]. It currently implements the ‘Union Rule Configuration’ (URC) rule evaluation scheme of Coomb’s et al [8,36] and an optimized version of the ‘Exhaustive search’ algorithm of Sokhansaj et al [**?**]. The Utilities Module consists of two submodules: a) the Postprocessing Submodule and, b) the Add-ons (or Plug-ins) Submodule. The Postprocessing Submodule consists of three Units – the ‘Graph’, ‘Evaluation (or Validation)’, and the ‘Dynamic Simulation’ Units. The Graph Unit consolidates the best fitted models derived from the fuzzy inference system into a network graph as explained in Gormley et al [15]. The Evaluation Unit simply compares expression profile predictions by inferred models to the experiment observed values. Depending on the user-specification, this may be against original model elucidation data (default) or an independent dataset. Also depending on the user, a re-calculation of the model’s fit may be specified, particularly to quantitatively describe how well models fit independent datasets. The ‘Dynamic Simulation Unit’ implements and executes model dynamic simulations as also described in Gormley et al [15]. To facilitate downstream data integration, the dynamic simulations’ stop criteria is dependent on the user-specified number of iteration steps and not the computed error. However the error estimates at the end of the simulation runs are reported. In anticipation of community contributions, the Add-on (or Plug-ins) Submodule is described. Current in-house created functionalities that would fit appropriately configured add-on units include an “In-Silico Knockout Simulations” add-on and a “Visualization” add-on which depends on a secondary-installed program. These are also freely available on request.

**Figure 1:**
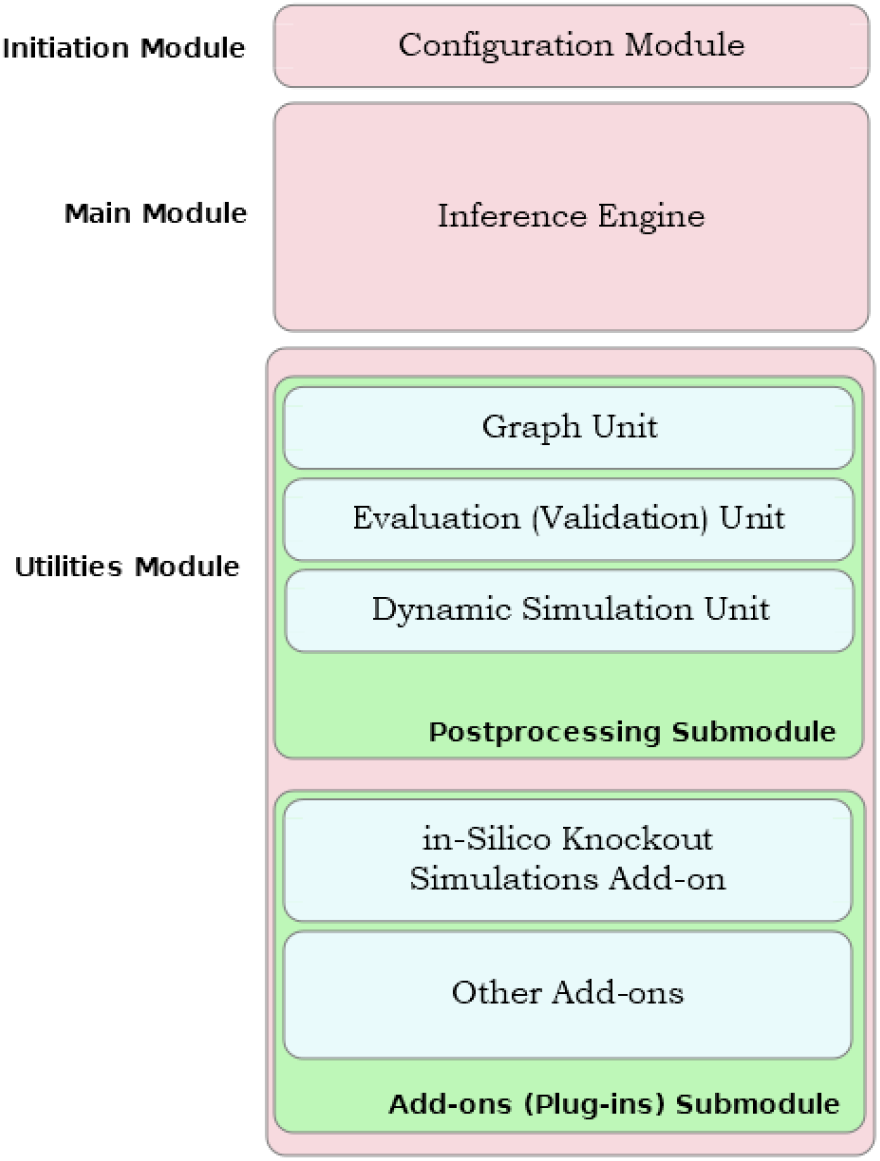
The jFuzzyMachine Application Components

### 2.4 Getting jFuzzyMachine

`jFuzzyMachine`’s source codes and precompiled binaries are freely available at the Github repository locations https://github.com/paiyetan/jfuzzymachine and https://github.com/paiyetan/jfuzzymachine/releases/tag/v1.7.21 respectively.

### 2.5 Installation Requirements

`jFuzzyMachine` is platform independent. It would run on a Windows, Mac, or UNIX-based Operating System (OS) with an appropriately preinstalled Java Runtime Environment (JRE). Java 7 or above is required. You may download the latest version of Java from https://www.java.com/en/download/.

To run the visualization add-on (plugin), provided as an added-value, a UNIX-based OS with the R program statistical computing environment preinstalled, is required. R may be downloaded from https://cran.r-project.org/.

### 2.6 Installing jFuzzyMachine

Unzip the compressed application package into a directory of choice. The content of the unzipped folder should include: One primary java archive (.jar) folder, four runtime configuration (.config) files, and four subdirectories (`etc/, lib/, plugins/`, and `src/`),

- `JFuzzyMachine.jar`
- `jfuzzymachine.config`
- `jfuzzymachine.graph.config`
- `jfuzzymachine.evaluator.config`
- `jfuzzymachine.simulator.config`
- `etc`
- `lib`
- `plugins`
- `src`

The configuration files are pre-filled to satisfy required parameters for this manual’s demonstration. Users may appropriately fill-in their own specifications and experiment with the tool. See configuration options below.

### 2.7 Running jFuzzyMachine

To run the tool, on the command-line,

1. Navigate into the application directory
2. Appropriately fill-in the desired run-time options in the configuration files and
3. Depending on application module or functional unit of interest, type the following commands, one at a time:

To elucidate fuzzy logic-based regulatory relationships, run the commands

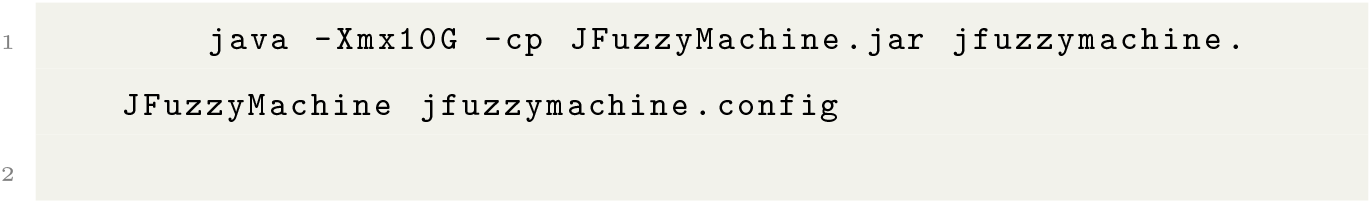

To derive a composite network graph, including rule frequencies, run

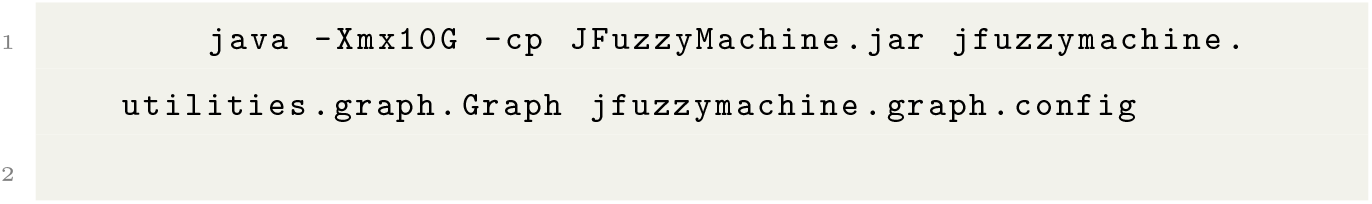

To evaluate or validate how well inferred fuzzy logic-based regulatory models fit the data, run

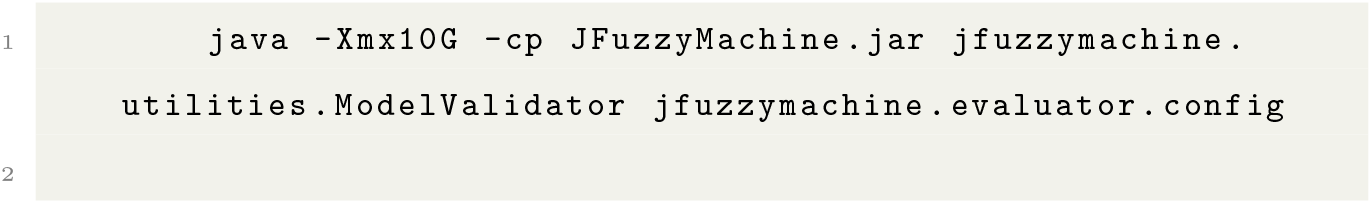

To run dynamic simulation of regulatory network, and tease expression values at systems steady state, run

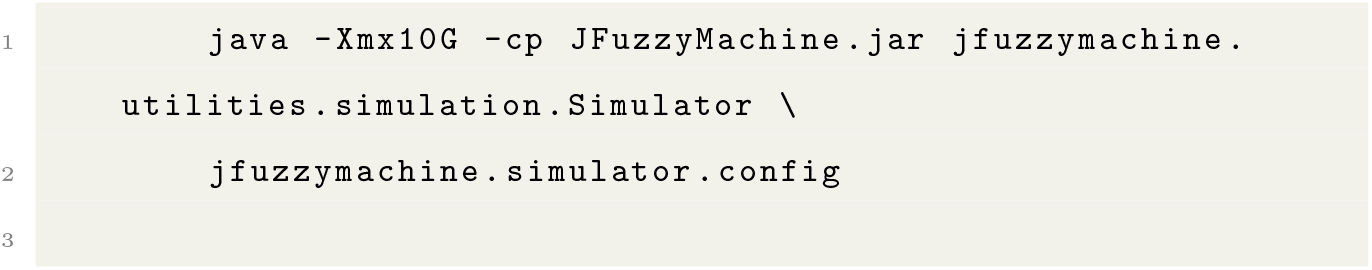

The `-Xmx10G` option indicates the amount of memory space available to the program during the program execution. It is specified as 10 Gigabytes. For very large expression data, it is recommended to be increased. The `-cp` option indicates the location of our Java program class files – specified as `JFuzzyMachine.jar`. It is assumed this is located in the current working director.

#### 2.7.1 The jfuzzymachine.config file

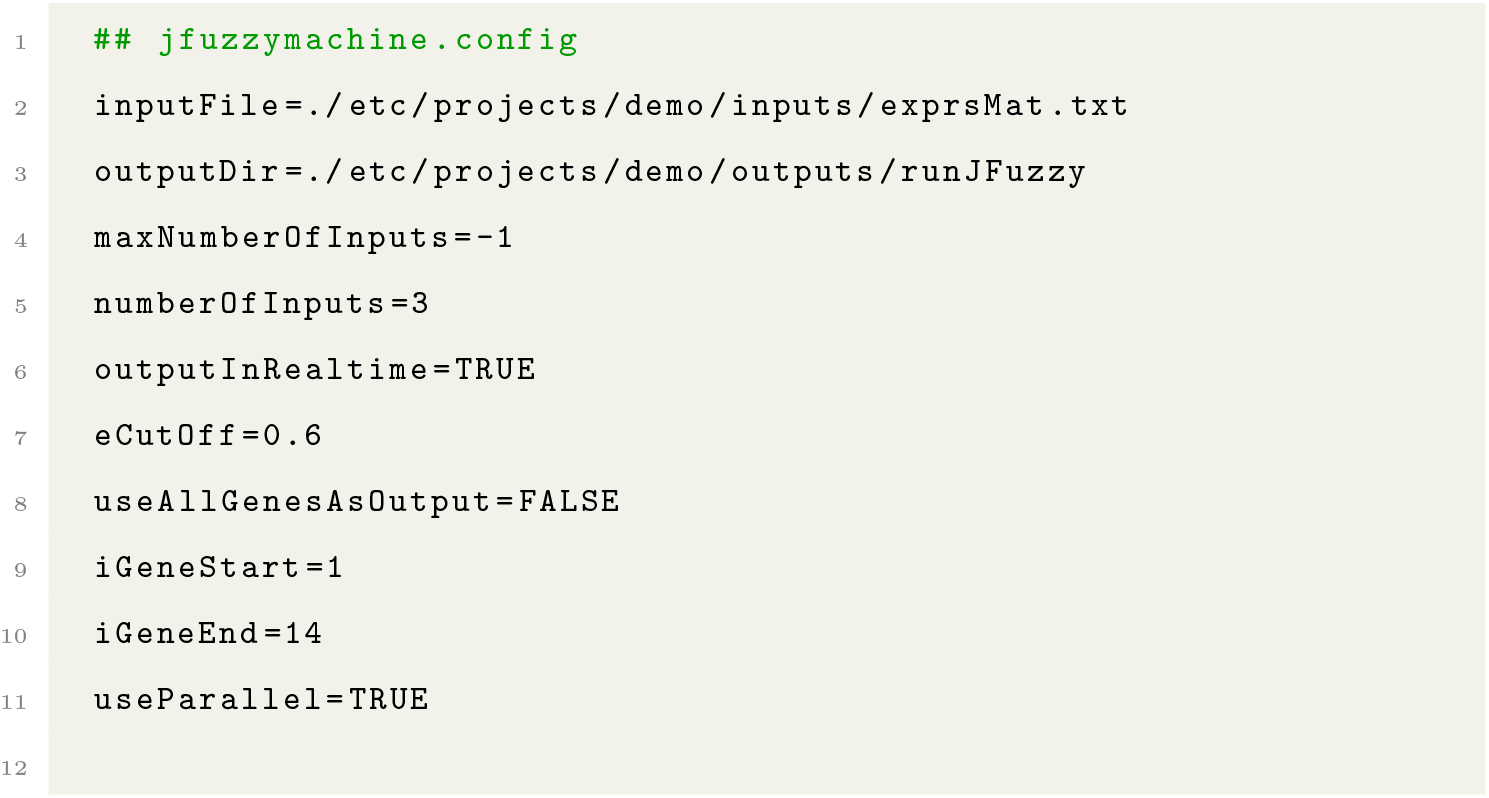

The `jfuzzymachine.config` has, at least, the above listed parameters (‘key’=‘value’ pairs). The associated values listed here are for demonstration purposes in this manual. The `inputFile` option specifies the relative path to the data matrix of normalized expression values. The `outputDir` specifies the path to the directory where results from `jFuzyMachine` are to be placed. The `maxNumberOfInputs` option is a flag that specifies how `jFuzzyMachine` should handle input (regulatory) features. A negative flag indicates that exactly the specified `numberOfInputs` option be considered. A positive value specifies to `jFuzzyMachine` to consider all possible number of inputs up-to the specified value. E.g. a positive value of 4, simply says to `jFuzzyMachine` to consider all possible combinations of 1, 2, 3, and 4 regulatory inputs to an output feature. Current implementation of `jFuzzyMachine` allows up to 5 inputs. A negative flag however, says to `jFuzzyMachine` to consider only possible combinations of 3 regulatory inputs (specified by the `numberOfInputs` option in the above configuration) to an output feature. The `outputInRealtime` option tells `jFuzzyMachine` to output its runtime informations onto a standard output (the console). This typically includes derived models, inferred regulatory rules, and computed fit estimates. The `eCutOff` option specifies the cut-off for which to consider a computed fuzzy logic model. Models below the specified value are discarded. The `useAllGenesAsOutputs` option specifies whether to consider all features in the expression values matrix or a limited set specified by the `iGeneStart` and `iGeneEnd` options. The `iGeneStart` and `iGeneEnd` options specify the range of features to use from a numerically ordered list – the expression matrix row numbers. The options `iGeneStart=1` and `iGeneEnd=14` in the configuration above, simply says to `jFuzzyMachine` to consider features of expression profiles from the first to the 14^th^ row, in the expression matrix (`inputFile`), as probable outputs in the regulatory model inference. The `useParallel` option indicates whether to run `jFuzzyMachine` in the optimized mode for speed (distributed across computing cores available at runtime).

#### 2.7.2 The jfuzzymachine.graph.config file

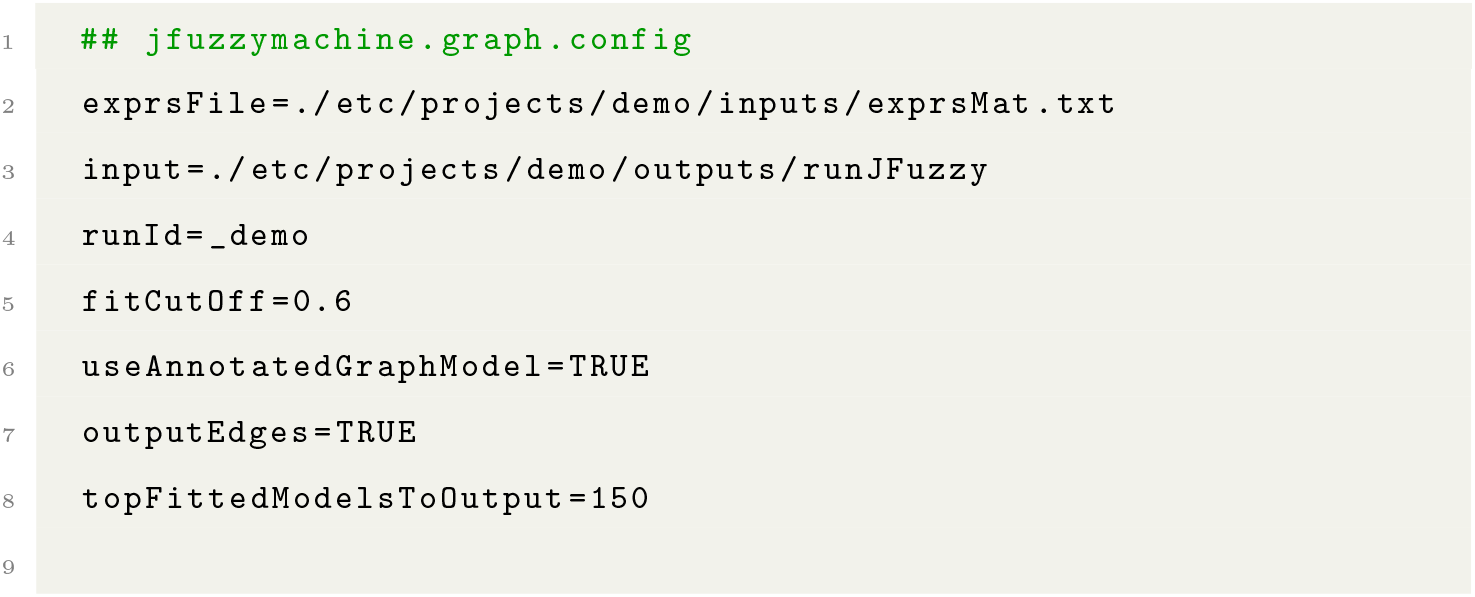

The above are parameters (runtime options) to the `jFuzzyMachine` Graphical unit. The `exprsFile` option specifies a path to the expression matrix from which regulatory models were inferred. The `input` option specifies a path to the directory in which `jFuzzyMachine` inferred models and output result files are placed. The `runId` option is a user-specified identifier prepended to outputted results’ file-names (please see the Results section). The nomenclature (naming convention) of the outputted result files is of the form *<*`runId`> `runJFuzzUtils.`*<*`fileType`>. The `fitCutOff` option specifies a cut-off for considering models. Models above specified `fitCutOff` are considered for inclusion in a consolidated network model. Only the regulatory edges of the passing models are considered. The `useAnnotatedGraphModel` tells `jFuzzyMachine` Graph unit to model regulatory network as a directed acyclic graph. If `TRUE`, the outputted adjacency matrix (`.adj` or `.mat` file) is a directed graph. By default, `jFuzzyMachine’s` Graphical unit outputs only an adjacency matrix file, to represent the inferred regulatory network, but the option `outputEdges` specifies to `jFuzzyMachine` to also print edges (a `.edg` file) of the consolidated network. The `topFittedModelsToOutput` applies to ouputting fitted models. It specifies the number of alternate top ranked models that pass `fitCutOff` filter option to report in the output (`.fit2`) file.

#### 2.7.3 The jfuzzymachine.evaluator.config file

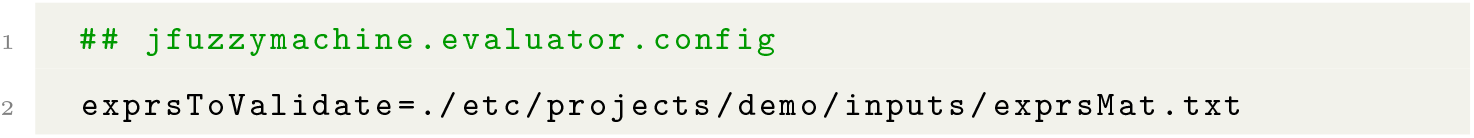
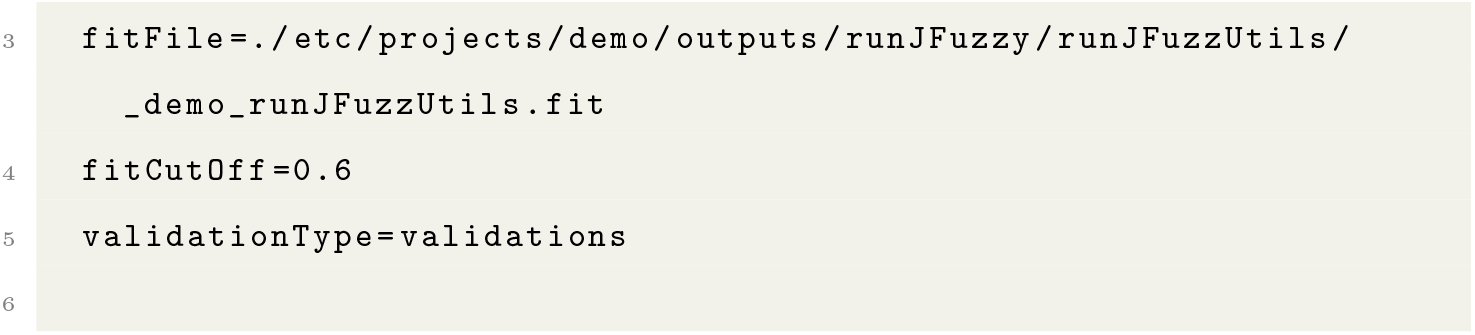

In `jfuzzymachine.evaluator.config` file, the `exprsToValidate` option specifies a path to the expression matrix from which regulatory models were inferred – in the case of evaluating the performance of the inferred models against the model– generating data. For an independent evaluation of the model, this is a path to the expression matrix of the independent dataset. The `fitFile` specifies the path to derived `.fit` file from j`FuzzyMachine’s` Graph unit. The `.fit` file contains the best fitted models. The `fitCutOff` specifies an estimated fit cut–off value above which to consider models. The `validationType` option, specifies what validation is being performed. Acceptable values include `validations` (default) and `ivalidations`. Specifying `ivalidations` implies an independent validation is being performed.

#### 2.7.4 The jfuzzymachine.simulator.config file

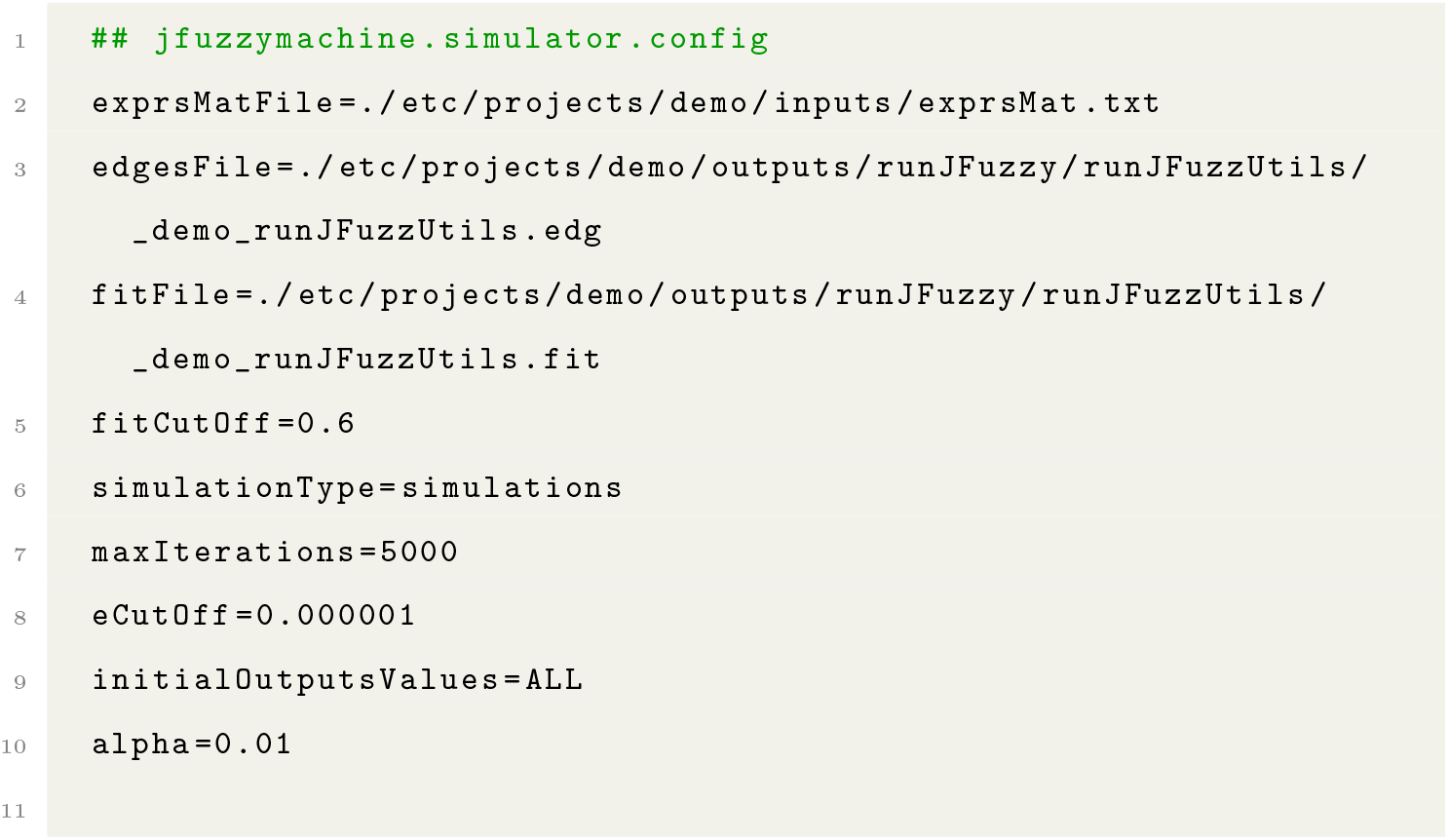

In the `jfuzzymachine.simulator.config`, the `exprsMatFile` specifies a path to the expression matrix from which regulatory models were inferred – in the case of evaluating the performance of the inferred models against the model–generating data. For an independent evaluation of the model, this is a path to the expression matrix of the independent dataset. The `edgesFile` and `fitFile` options specify the path to the `.edg` and `.fit` files derived from `jFuzzyMachine’s` Graph unit. These contain the edges of the consolidated network and the best fitted models from the regulatory model elucidating steps. The `fitCutOff` specifies an estimated fit cut-off value above which to consider models. The `simulationType` option specifies the sort of dynamic simulation to be performed. Acceptable values include `simulations` (default) and `isimulations`. Specifying `isimulations` implies a dynamic simulation of consolidated network, using previously derived models as simulation parameters on an independent dataset is being performed. The `maxIterations, eCutOff, initialOutputsValues` and `alpha` are other dynamic simulation parameters. The `maxIteration` and `eCutOff` are the stopping criteria – maximum iteration steps and error estimate cut–off respectively. Default values are 5000 and 10*e* ^*-*7^ respectively. To better facilitate integration with downstream analyses, the Dynamic Simulation Unit defaults to preferably using the `maxIteration` option as stopping criteria. The `initialOutputsValues` specifies which ‘perturbation’, ‘sample’, or ‘time–point’ values, from the expression matrix, to use as initial values in the simulation. It defaults to `ALL`, i.e. all values are sequentially used. Other values are `FIRST` and `RANDOM`, implying the first column and a random column values respectively. The `alpha` option specifies the ‘mixing parameter’, *α*, of the simulation model. Based on Gormley et al, linear combination of new and old values ensures that the system smoothly converges towards equilibrium. And accordingly, `jFuzzyMachine`’s Dynamic Simulation Unit computes new values of each node (*I*_1_) based on the initial conditions and the fuzzy relations inferred from the data; values in the next iteration were calculated as a linear combination of the inferred values (*I*_*n*_) and the initial values (*I*_*n-*1_) as follows:

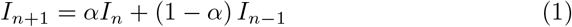

### 2.8 Comparing jFuzzyMachine’s inferred network to ARACNe’s

To compare regulatory network inferred by `jFuzzyMachine` with that inferred by the ARACNe (an Algorithm for the Reconstruction of Gene Regulatory Networks algorithm [26], mutual information matrix of sampled features expression profile was inferred using the R/bioconductor `minet` package `build.mim` routine and specifying the `spearman` option as the estimator. The `minet` package ARACNe algorithm implementation was used to derived weighted adjacency matrix of the inferred network. The identified edges were compared to those inferred from the best fitted models from a `jFuzzyMachine` inference, given the same expression profile.

## 3 RESULTS AND DISCUSSION

In this section we describe results outputted from a typical `jFuzzyMachine` execution. These are are described as results produced by the different modules of the tool - the Main module and Utilities module. Presented results are those of the sampled data expression profile.

In the second part, we describe, the results obtained from a comparison of the results of `jFuzzyMachine` and that of the `ARACNe` algorithm with respect to potential regulatory interactions derived from the sampled data.

### 3.1 The Main Module - Inference Engine

The main output results from the `jFuzzyMachine` inference engine are written to the `outputDir`. These are files or single file ending with `.jfuz`. From the demo run, this would be the `./etc/projects/demo/outputs/runJFuzzy/exprs-Mat.1.14.3.TRUE.jfuz`. This consists of 4 major sections indicated by the > character at the begining of the line. These sections include; a prologue, run parameter listing, the main result, and an epilogue. The prologue section stores information such as the run’s start-time, while the epilogue stores the run end-time and duration. The main section is a tab-delimited table with columns: `Output, NumberOfInput(s), Input(s), Rule(s)`, and `Error(E)`. The `Output` column indicates the output node in the model; the `NumberOfInput` indicate the number of input nodes, the `Input(s)`, considered. The `Rule(s)` column indicate the fuzzy logic rule that associates the respective input node to the output node. The `Error(E)` column indicates the model’s fit. Shown below is a sample output from the demo run:

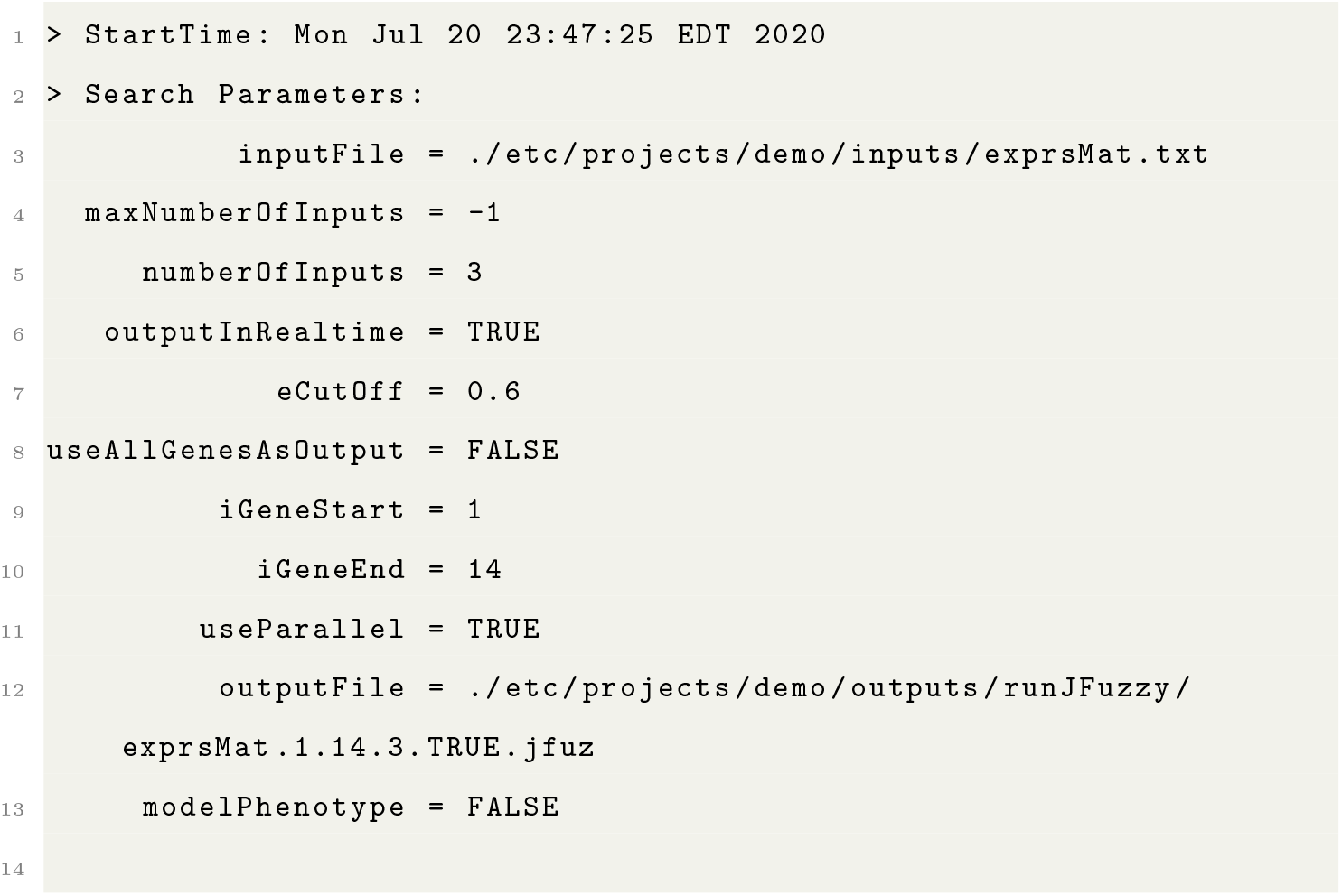
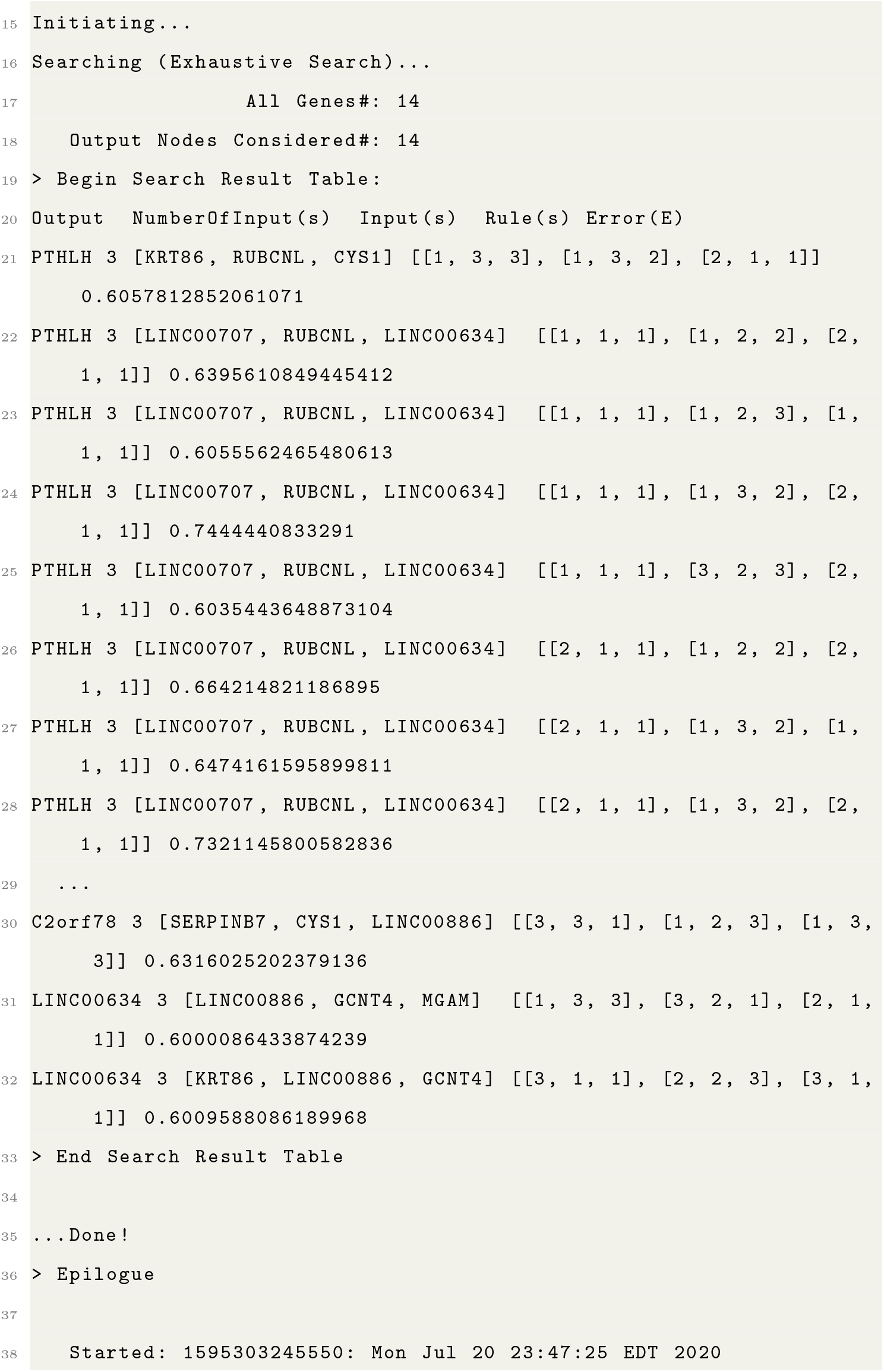
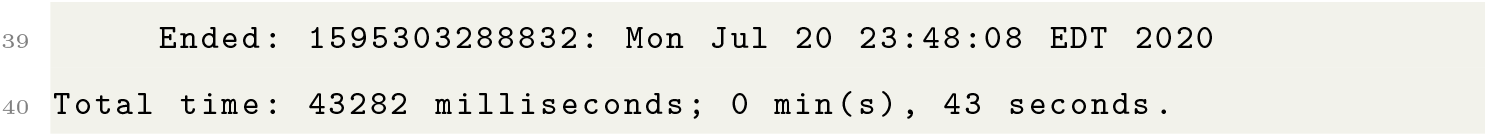

### 3.2 The Utilities Module

`jFuzzyMachine’s` Utilities Module consists of the ‘Postprocessing’ and the ‘Addons’ submodules. The postprocessing module consists of the ‘Graph’, ‘Evaluation’ and ‘Dynamic Simulations’ Units.

#### 3.2.1 The Graph Unit

Outputs from the graph unit execution are placed in the `runJFuzzUtils` subdirectory. With regards to this demonstration, this would be the `./etc/projects/demo/outputs/runJFuzzy/runJFuzzUtils` directory. These tab-delimited result files include:

- `_demo_runJFuzzUtils.adj`
- `_demo_runJFuzzUtils.edg`
- `_demo_runJFuzzUtils.edg2`
- `_demo_runJFuzzUtils.fit`
- `_demo_runJFuzzUtils.fit2`
- `_demo_runJFuzzUtils.fre`

The `_demo_runJFuzzUtils.adj` file is a directed graph adjacency matrix describing the connections in the inferred network. A connection between two nodes is indicated by 1 and 0 vice versa. Features in the rows are the inputs while those in columns are the output nodes. The `_demo_runJFuzzUtils.edg` and `_demo_runJFuzzUtils.edg2` result files are about the same. Describing the edges in the inferred network, they both have the columns; `From, To, Rule`, and `Weight` in common. These correspond to the input node, output node, fuzzy logic rule associating the input with the output node, and estimated model fit respectively. The ‘HashCode’ column in the “.edg” file is only included for programmatic debugging. Likewise, the `_demo_runJFuzzUtils.fit` and `_demo_runJFuzzUtils.fit2` result files are about the same. While the `.fit2` reports all models above the specified `fitCutOff` in the `jfuzzymachine.graph.config` file, the `.fit` file reports only the best fitted model to each output node. Sampled outputs from the related demo run are shown below:

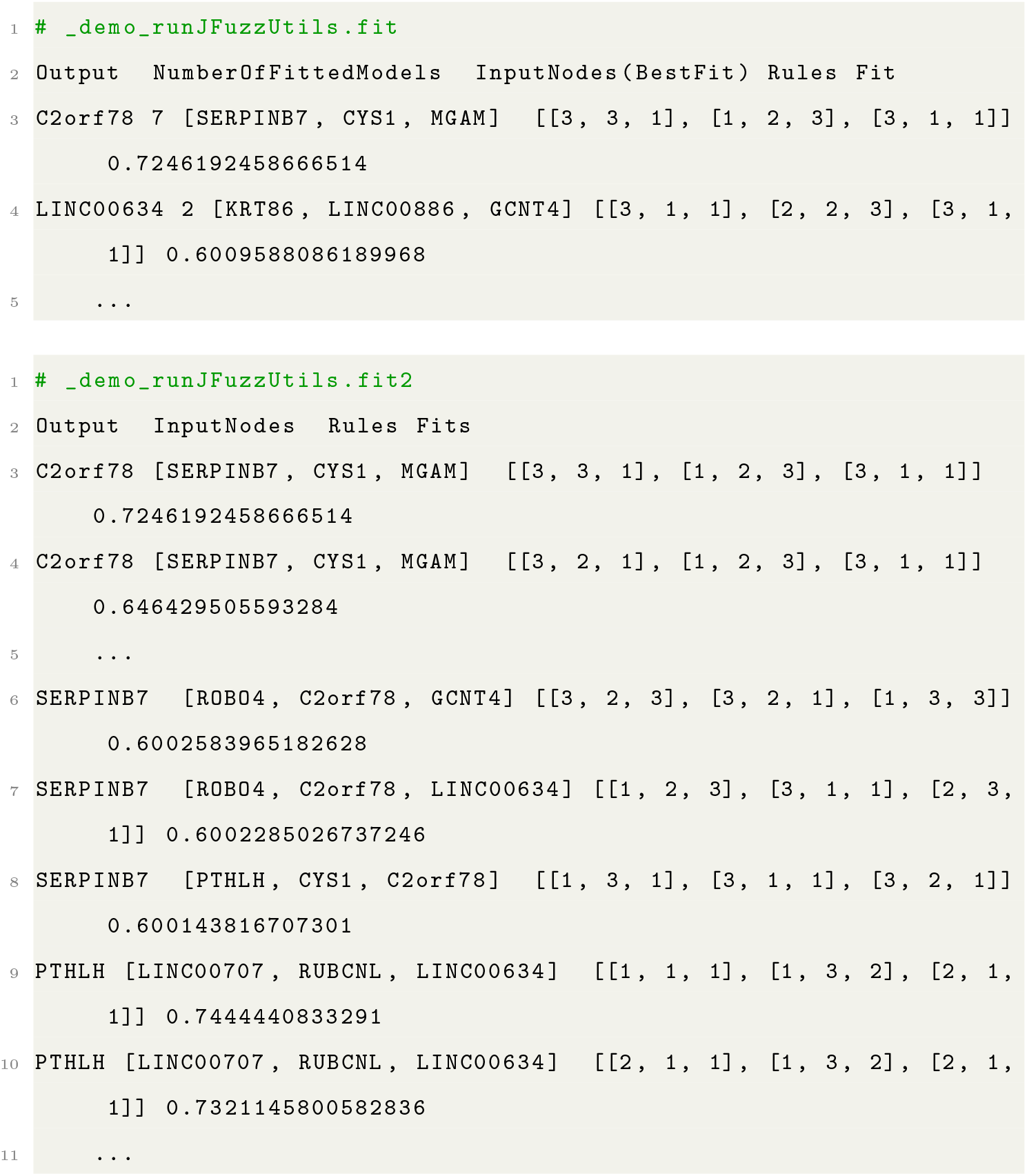

The `_demo_runJFuzzUtils.fre` reports the frequency of the fuzzy rules evaluated in the inferred models with an estimated fit value above the `fitCutOff`. Please see Gormley et al and Sokhansanj et al for a detailed explanation of the rules.

#### 3.2.2 The Evaluation Unit

The Evaluation Unit compares expression profile predictions by inferred models to an experiment values – either the fuzzy logic models’ model-elucidating data or an independendent dataset. Its output are reported in the `_demo_runJFuzzUtils.val` file – an expression matrix of predicted values of output nodes, given the values in the `exprsToValidate` file and the set of fuzzy logic models in the `fitFile` specified in the `jfuzzymachine.evaluator.config`.

#### 3.2.3 The Dynamic Simulation Unit

The ‘Dynamic Simulation Unit’ implements and executes model dynamic simulations as also described in Gormley et al. The unit implements an iterative scheme to determine the state of the network at equilibrium. Simulation values are reported in the `runJFuzzyUtils/simulations/` subsub-directory in the `jFuzzyMachine` main output directory. These are captured in the `.dta` and `.sim` files. The `.dta` files report the error values following each iteration, while the `.sim` file reports the estimate for each output node in the network at each iteration. The numerical value in the naming convention of the derived files show the column index, in the expression matrix, of the ‘sample’, ‘perturbation’, or ‘time-point’ from which initial values for the respective simulations were derived.

### 3.3 Add-ons

`jFuzzyMachine` and its outputs are designed to either be standalone resources, or be easy to integrate with other analyses pipelines and platforms. Add-ons or plug-ins provide an avenue to easily integrate additional functionalities or integrate other tools to the base application. For a better appreciation of results, we have included an example plug-in to enable some visualization of results demonstrated in this manual. As previously stated (please see publication), plug-ins can be platform dependent and may rely on secondary applications for full functionality. The plug-in bundled with `jFuzzyMachine` requires a UNIX-based platform or OS with the R statistical programming environment pre-installed. In addition to having the R program pre-installed, the following R/Bioconductor packages are required:

- `optparse`
- `org.Hs.eg.db`
- `xtable`
- `igraph`
- `graph`
- `Rgraphviz`
- `pheatmap`
- `ReactomePA`

To execute, simply run the following commands from within the `jFuzzyMachine` application working directory:

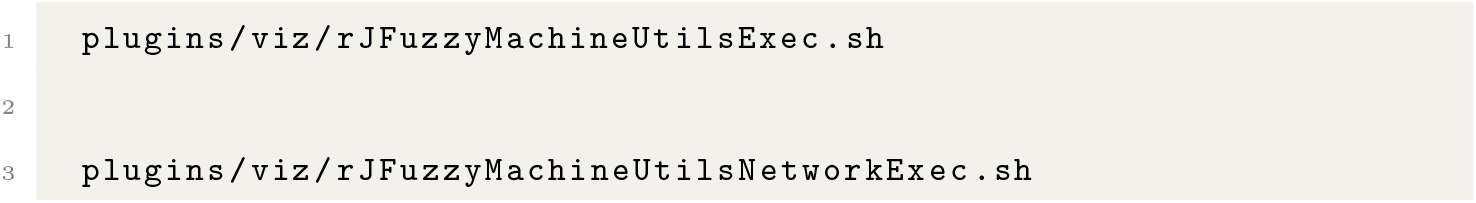

Example output figures, saved in the `./etc/projects/demo/outputs/plugins/viz/figs` directory are presented below:

Figure 2 shows the predicted values by the best fitted models for a the sample output node, the C2orf78 gene product. It can be appreciated that predicted values across the assay samples tend to trend similarly as the observed values indicated in gray. `jFuzzyMachine` predictions are model-based. A model consists of a set on input nodes and an output node with regulatory rule relationships among these.

**Figure 2:**
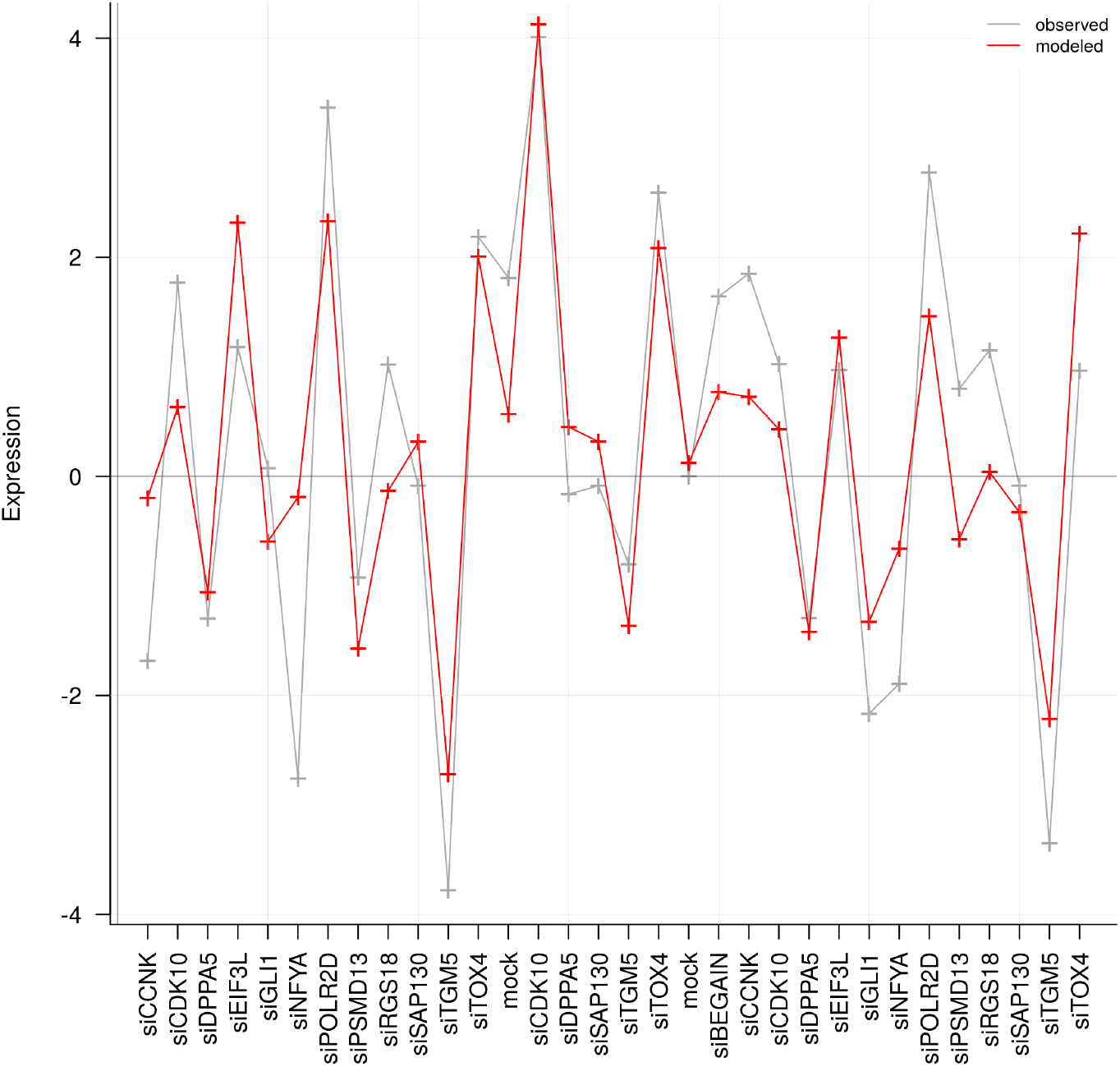
An Evaluation (Validation) Plot. A visual evaluation of predictions of a fitted model for a sample output node, the **C2orf78** gene. The estimated fit was 0.72. The input (regulatory) nodes were the genes *SERPINB7, CYS1*, and *MGAM*. The y-axis indicates the normalized expression values and the a-axis indicates the sample perturbations or treatment. Samples were reverse transfected vorinostat-resistant colon cancer, HCT-116, cell lines. Each sample was treated with the indicated small interferring ribonucleic acid (siRNA) to knockdown the respectively indicated gene products. The grey plot line shows the observed expression profile of the gene **C2orf78**, while the “red” line shows the predicted expression value from the expression of the regulators in the given data, and the rules associating the regulators to the output. The inferred patterns of regulation (rules) are indicated in Figure 4. It can apparently be appreciated that the fuzzy logic model is able to tease out trend in the dataset.

Figure 3 shows predictions at steady states of an inferred regulatory network. Observed convergence gives a bit of added assurance that inferred relationships are likely to be feasible. A lack of convergence may indicate a suboptimal network inference secondary to suboptimal fits of derived models.

**Figure 3:**
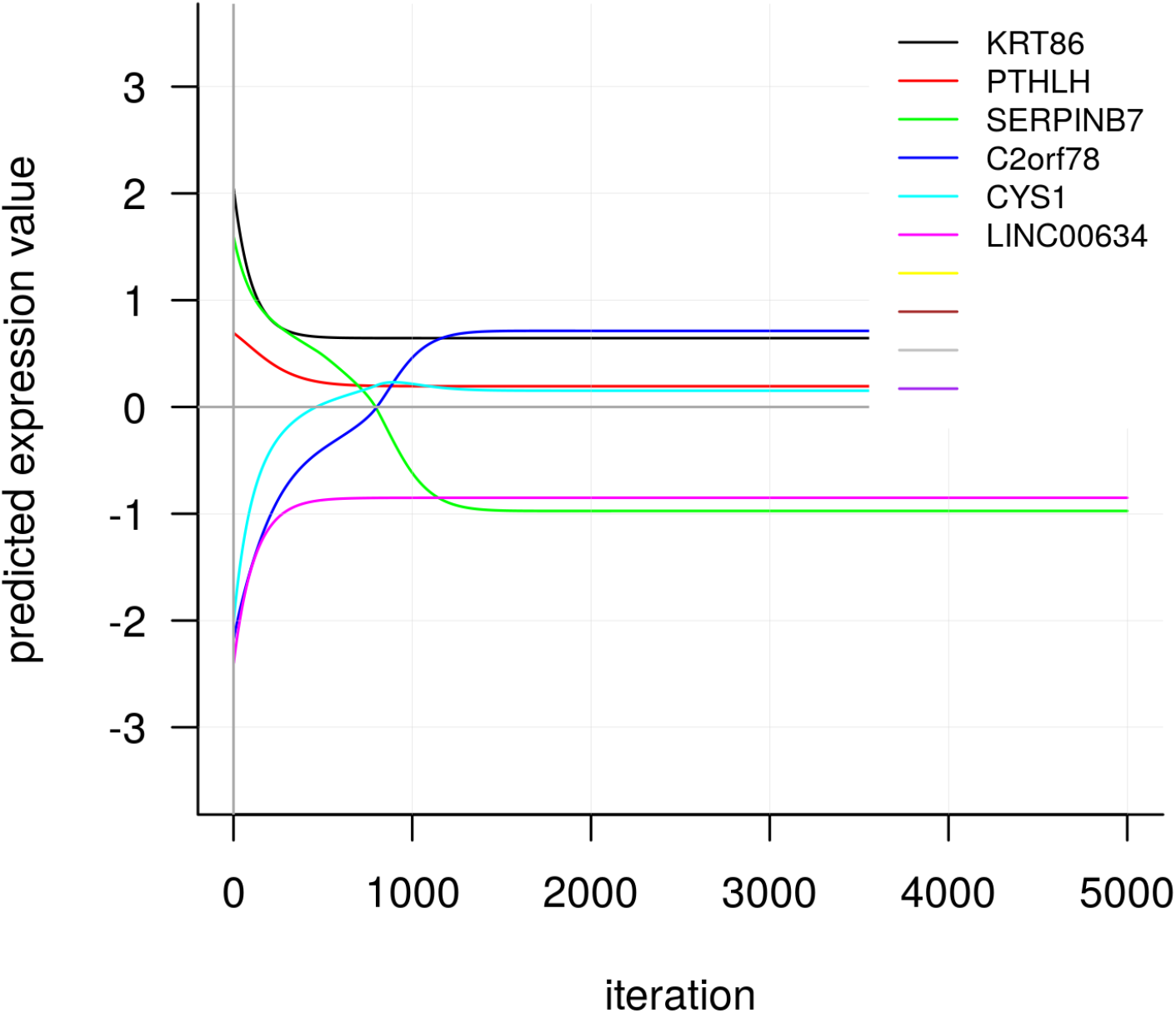
A Dynamic Simulation Plot. Having randomly chosen a sample (the 25th column sample) from the normalized expression matrix to provide initial values of expression, and given the best fitted models, the plot shows predicted expression values for the inferred outputs *KRT86, PTHLH, SERPINB7, C2orf78, CYS1* and *LINC00634* over 5000 iterations. It is appreciable that the inferred network achieves an equilibrium state at a little over 1000 iterations, when a change in predicted values tend to zero.

Figures 4 and 5 show the inferred networks of the `jFuzzyMachine` and that of the ARACNe algorithm respectively. It is observed that `jFuzzyMachine` both appear to have a large overlap in the number of predicted edge relationships. Fig. 6. `jFuzzyMachine` almost always predict the direction of relationship (i.e. what is regulating what). Given a evidences from the two tools of potential interaction between pairs of features, the biological significance of such interactions **may be worth further pursuit**.

**Figure 4:**
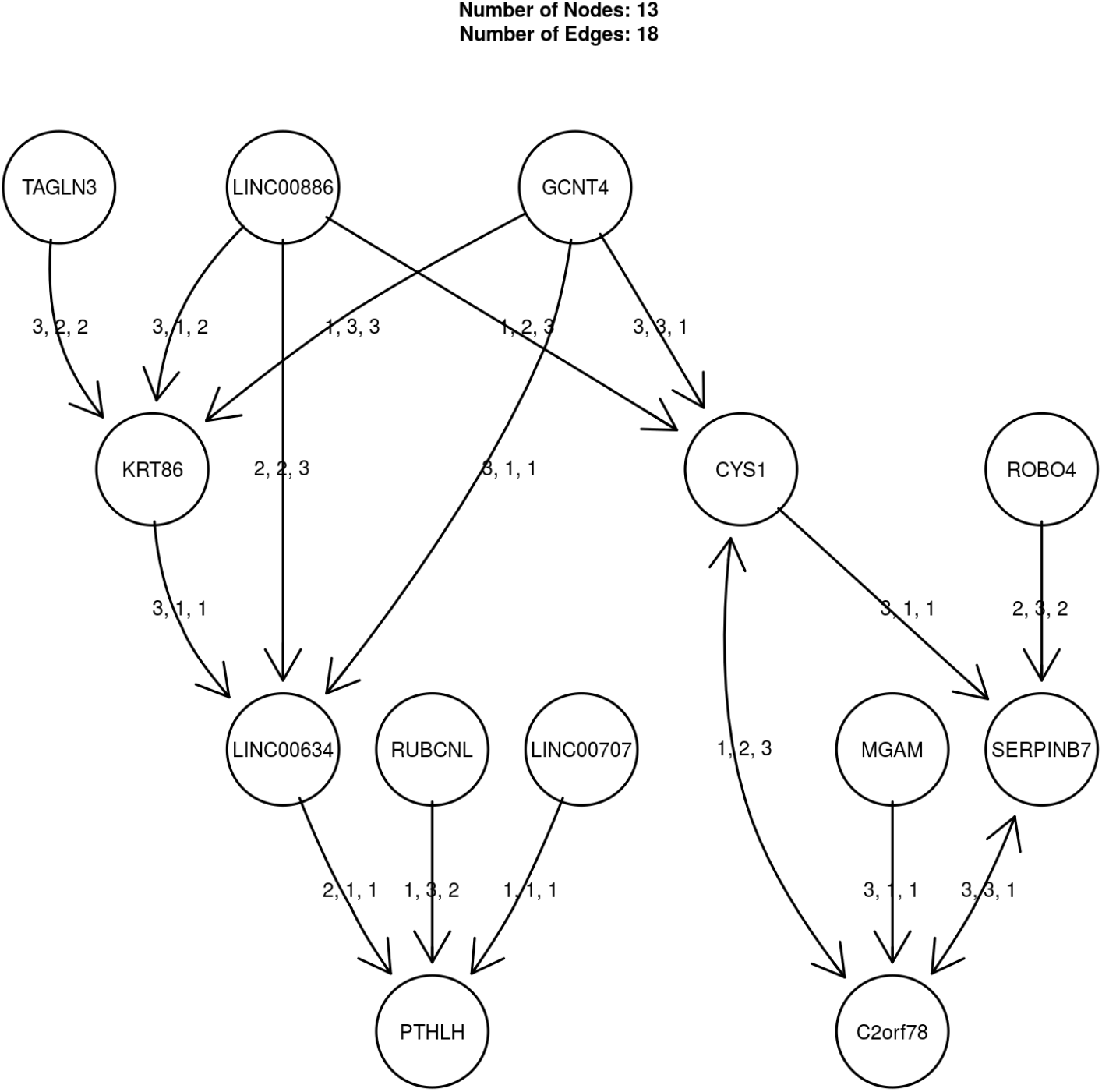
The Fuzzy Logic-based Regulatory Network Inferred. A composite regulatory network is inferred from the best fitted models for each node. The inferred network consists of 13 nodes (genes), and 18 edges (regulatory connections). The arrow heads indicate the regulatory direction from the input node to the output node. The edge labels, shown by the fuzzy rules, indicate the regulatory interaction. From Gormley et al, Rule configuration is the specification of if-then relationships between variables in fuzzy space. For example, an inhibitory relationship is represented by the rule vector *r* = [*r*_1_, *r*_2_, *r*_3_] = [3, 2, 1] (i.e., if input is low (*r*_1_), then output is high (3); if input is medium (*r*_2_), then output is medium (2), and if input is high (*r*_3_), then ouput is low (1). From the composite regulatory network, the regulatory effect of the *MGAM* gene on the *C2orf78* gene is indicated by the rule 3, 1, 1. This implies that when *MGAM* is low (*r*_1_), *C2orf78* is high (3); when it is medium (*r*_2_), *C2orf78* is low (1); and when *MGM* is high (*r*_3_), *C2orf78* is low (1). Notice that the bi-directional relationship between the pair of genes *C2orf78* –*CYS1*, and *C2orf78* –*SERPINB7*

**Figure 5:**
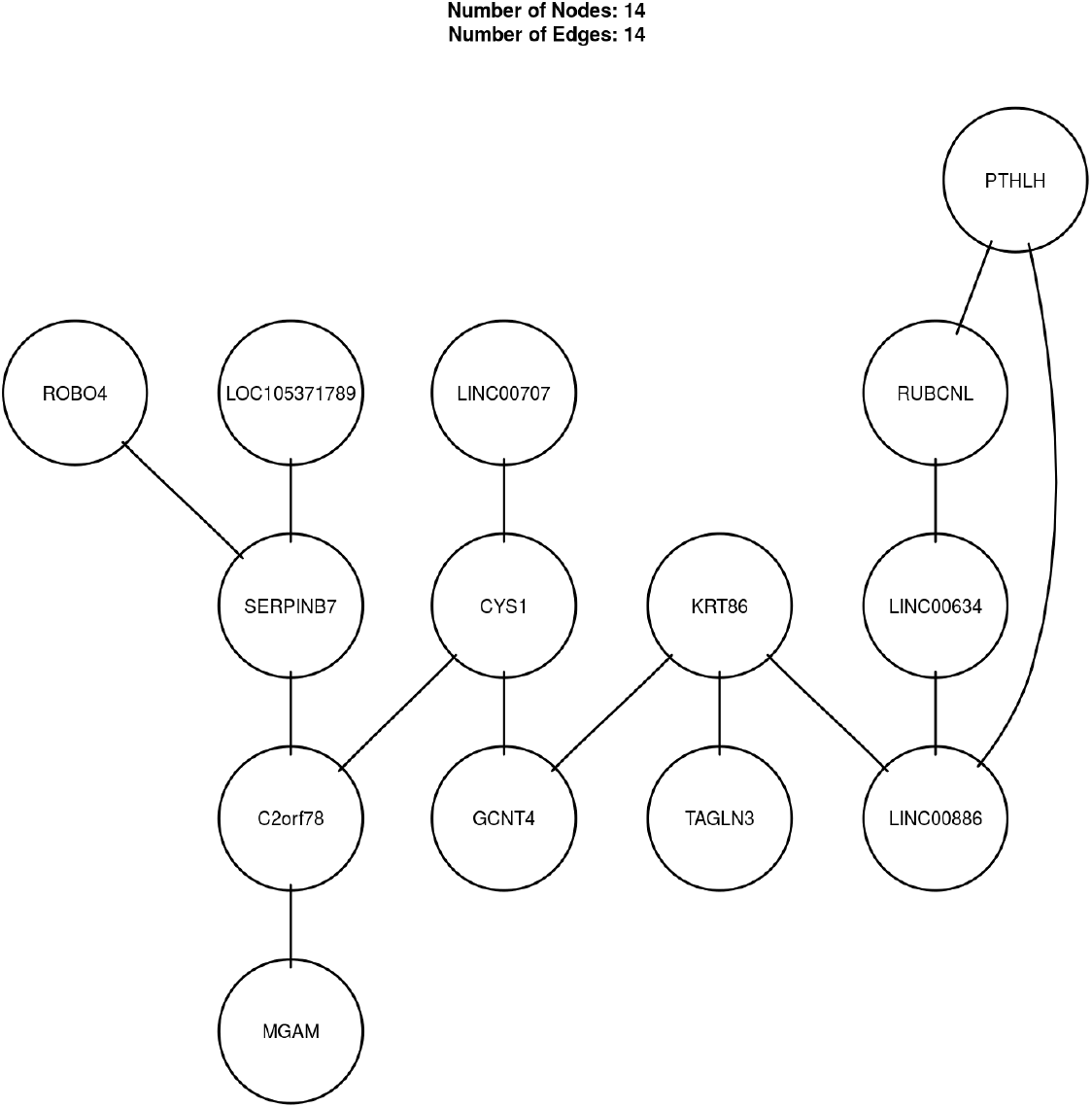
The ARACNe-inferred Regulatory Network. As described in the Materials and Methods section, mutual information between pairs of features was estimated using the correlation approach, to derive a mutual information matrix.

**Figure 6:**
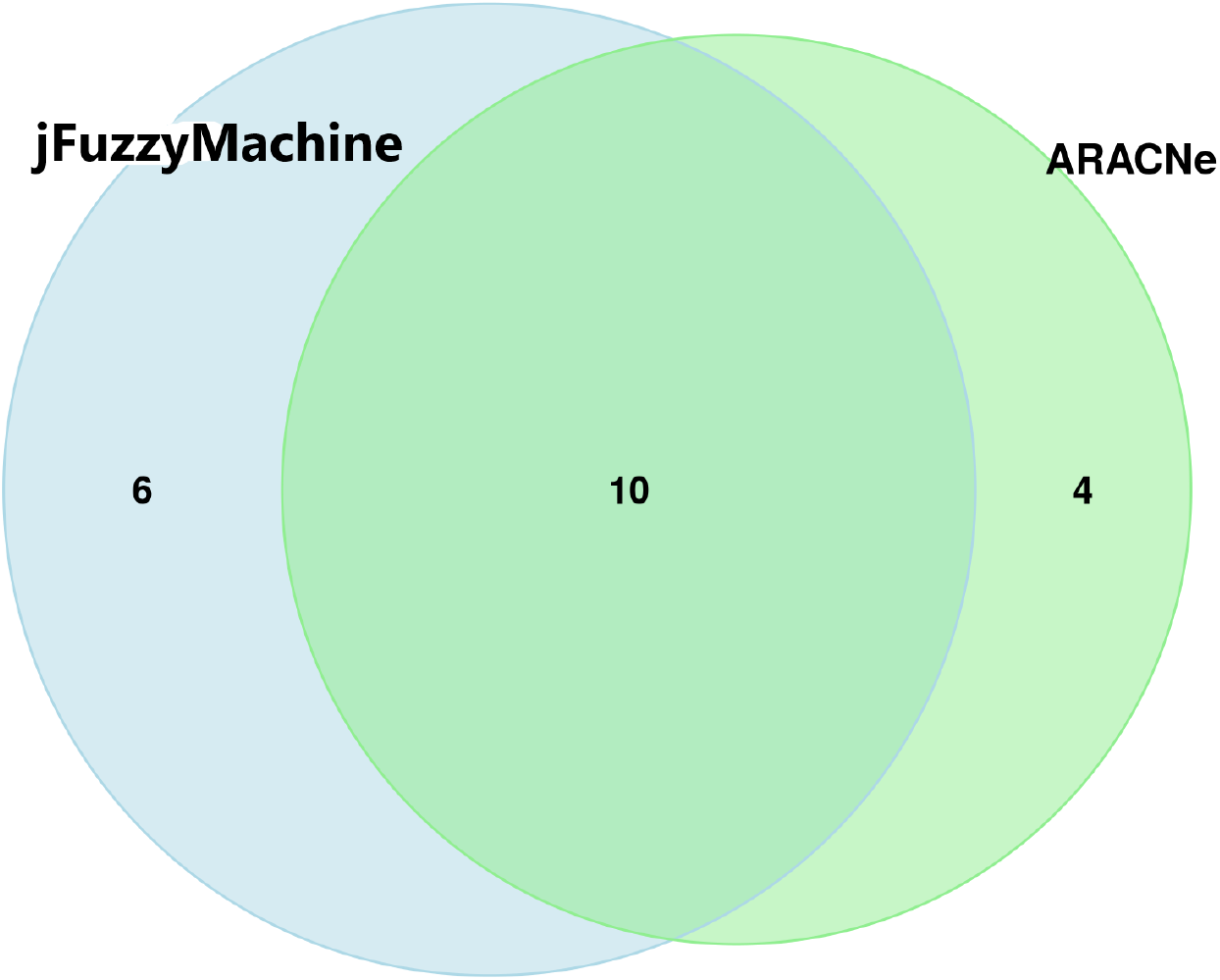
Identified edges between `jFuzzyMachine` and the `ARACNe` algorithm. A total of 16 edge relationships were identified by `jFuzzyMachine`. Two of these are observed to be by directional (Fig 2). The ARACNe algorithm identifies 14 network edges, over 70 percent of which are equally identified by `jFuzzyMachine`

## 4 CONCLUSION AND RECOMMENDATION

The Fuzzy logic inference approach to elucidating regulatory networks, although relatively mature, has little to no freely available tool to facilitate its adoption, to a greater extent, in elucidating high-throughput biological expression data. The `jFuzzyMachine` tool fills this apparent need. `jFuzzyMachine` mitigates the limitation of applying and appropriating the benefits of the fuzzy inference system to biological data. With respect to predicting regulatory interactions, `jFuzzyMachine` compares favorably with other regulatory inference tools. We expect to continue to include added functionalities and improve its current implementation. With our modular design and plan to accommodate third-party add-ons, we hope to facilitate community contributions and a scientific ecosystem of adopters. Also, we hope to continue to facilitate its integration with other high-throughput biological inference tools.

## Supporting information

jFuzzyMachine-Manual

## 5 ACKNOWLEDGEMENT

A special thanks to Dr. Iosif Vaisman, Dr. Dmitri Klimov, Dr. Saleet Jafri, and Dr. Quong at the George Mason University School of Systems Biology, for mentoring a larger body of the work for which a component part is presented here. The content presented here is solely the responsibility of the author and does not reflect the official views of any affiliated institutions.

## 6 AVAILABILITY

The jFuzzyMachine source codes and binaries are freely available at the Github repository locations: https://github.com/paiyetan/jfuzzymachine and https://github.com/paiyetan/jfuzzymachine/releases/tag/v1.7.21.

