## Supplementary material for "jFuzzyMachine – An Open–source Fuzzy Logic–based Regulatory Inference Engine for High–throughput Biological Data": jFuzzyMachine-Manual

### MANUAL
### jFuzzyMachine

Paul Aiyetan*[1,2] and Andrew Quong†[3]

[1]Frederick National Laboratory for Cancer Research, Frederick, MD 21701
[2]George Mason University School of Systems Biology, Manassas, VA 20110
[3]Fluidigm Corporation, San Francisco, CA 94080 (Current Affiliation)

#### Contents

**Introduction** **3**

**Getting jFuzzyMachine** **3**

**Installation Requirements** **3**

**Installing jFuzzyMachine** **3**

**Running jFuzzyMachine** **4**

**Add-ons** **12**

*
†

### Introduction

jFuzzyMachine is a fuzzy logic-based inference engine for biological high-throughput data. It addresses an apparent lack of readily available community tools implementing the Fuzzy Inference Systems with regards to high-throughput biological data. Please see publication.

### Getting jFuzzyMachine

jFuzzyMachine's source codes and precompiled binaries may be requested or freely downloaded from the bitbucket git repository locations https://bitbucket.org/paiyetan/jfuzzymachine/src/master/ and https://bitbucket.org/paiyetan/jfuzzymachine/downloads/. The application and distributed binaries are made available in a compressed folder named jFuzzyMachine.zip.

### Installation Requirements

jFuzzyMachine is platform independent. It would run on a Windows, Mac, or UNIX-based Operating System (OS) with an appropriately pre-installed Java Runtime Environment (JRE). Java 7 or above is required. You may download the latest version of Java from https://www.java.com/en/download/.

To run the visualization add-on (plugin), provided as an added-value, a UNIX-based OS with the R program statistical computing environment [1] pre-installed, is required. R may be downloaded from https://cran.r-project.org/.

### Installing jFuzzyMachine

Unzip the compressed application package into a directory of choice. The content of the unzipped folder should include: One primary java archive (.jar) folder, four runtime configuration (.config) files, and four subdirectories (etc/, lib/, plugins/, and src/),

- JFuzzyMachine.jar
- jfuzzymachine.config

[1] R Core Team (2019). R: A language and environment for statistical computing. R Foundation for Statistical Computing, Vienna, Austria. URL https://www.R-project.org/.

- jfuzzymachine.graph.config
- jfuzzymachine.evaluator.config
- jfuzzymachine.simulator.config
- etc
- lib
- plugins
- src

The configuration files are pre-filled to satisfy required parameters for this manual's demonstration. Users may appropriately fill-in their own specifications and experiment with the tool. See configuration options below.

#### Running jFuzzyMachine

To run the tool, on the command-line,

1. Navigate into the application directory
2. Appropriately fill-in the desired run-time options in the configuration files and
3. Depending on application module or functional unit of interest, type the following commands, one at a time:

   To elucidate fuzzy logic-based regulatory relationships, run the commands

```
       java -Xmx10G -cp JFuzzyMachine.jar jfuzzymachine.JFuzzyMachine \
                 jfuzzymachine.config

```

   To derive a composite network graph, including rule frequencies, run

```
       java -Xmx10G -cp JFuzzyMachine.jar jfuzzymachine.utilities.graph.Graph \
                 jfuzzymachine.graph.config

```

   To evaluate or validate how well inferred fuzzy logic-based regulatory models fit the data, run

```
      java -Xmx10G -cp JFuzzyMachine.jar jfuzzymachine.utilities.ModelValidator \
                jfuzzymachine.evaluator.config

```

To run dynamic simulation of regulatory network, and tease expression values at systems steady state, run

```
      java -Xmx10G -cp JFuzzyMachine.jar \
       jfuzzymachine.utilities.simulation.Simulator jfuzzymachine.simulator.config

```

##### The **jfuzzymachine.config** file

```
## jfuzzymachine.config
inputFile=./etc/projects/demo/inputs/exprsMat.txt
outputDir=./etc/projects/demo/outputs/runJFuzzy
maxNumberOfInputs=-1
numberOfInputs=3
outputInRealtime=TRUE
eCutOff=0.6
useAllGenesAsOutput=FALSE
iGeneStart=1
iGeneEnd=14
useParallel=TRUE

```

#### The **jfuzzymachine.graph.config** file

```
## jfuzzymachine.graph.config
exprsFile=./etc/projects/demo/inputs/exprsMat.txt
input=./etc/projects/demo/outputs/runJFuzzy
runId=_demo
fitCutOff=0.6
useAnnotatedGraphModel=TRUE
outputEdges=TRUE
topFittedModelsToOutput=150

```

The above are parameters (runtime options) to the jFuzzyMachine Graphical unit. The exprsFile option specifies a path to the expression matrix from which regulatory models were inferred. The input option specifies a path to the directory in which jFuzzyMachine inferred models and output result files are placed. The runId option is a user-specified identifier prepended to outputted results' filenames (please see the Results section). The nomenclature (naming convention) of the outputted result files is of the form <runId>\textsf{_runJFuzzUtils.}<fileType>. The fitCutOff option specifies a cut-off for considering models. Models above specified fitCutOff are considered for inclusion in a consolidated network model. Only the regulatory edges of the passing models are considered. The useAnnotatedGraphModel tells jFuzzyMachine Graph unit to model regulatory network as a directed acyclic graph. If TRUE, the outputted adjacency matrix (.adj or .mat file) is a directed graph. By default, jFuzzyMachine's Graphical unit outputs only an adjacency matrix file, to represent the inferred regulatory network, but the option outputEdges specifies to jFuzzyMachine to also print edges (a .edg file) of the consolidated network. The topFittedModelsToOutput applies to ouputting fitted models. It specifies the number of alternate top ranked models that pass fitCutOff filter option to report in the output (.fit2) file.

##### The **jfuzzymachine.evaluator.config** file

```
## jfuzzymachine.evaluator.config
exprsToValidate=./etc/projects/demo/inputs/exprsMat.txt
fitFile=./etc/projects/demo/outputs/runJFuzzy/runJFuzzUtils/_demo_runJFuzzUtils.fit
fitCutOff=0.6
validationType=validations

```

In jfuzzymachine.evaluator.config file, the exprsToValidate option specifies a path to the expression matrix from which regulatory models were inferred – in the case of evaluating the performance of the inferred models against the model–generating data. For an independent evaluation of the model, this is a path to the expression matrix of the independent dataset. The fitFile specifies the path to derived .fit file from jFuzzyMachine's Graph unit. The .fit file contains the best fitted models. The fitCutOff specifies an estimated fit cut–off value above which to consider models. The validationType option, specifies what validation is being performed. Acceptable values include validations (default) and ivalidations. Specifying ivalidations implies an independent validation is being performed.

##### The **jfuzzymachine.simulator.config** file

```
## jfuzzymachine.simulator.config
exprsMatFile=./etc/projects/demo/inputs/exprsMat.txt
edgesFile=./etc/projects/demo/outputs/runJFuzzy/runJFuzzUtils/_demo_runJFuzzUtils.edg
fitFile=./etc/projects/demo/outputs/runJFuzzy/runJFuzzUtils/_demo_runJFuzzUtils.fit
fitCutOff=0.6
simulationType=simulations
maxIterations=5000
eCutOff=0.000001
initialOutputsValues=ALL
alpha=0.01

```

[2] Gormley M, Akella VU, Quong JN, Quong AA. An integrated framework to model cellular phenotype as a component of biochemical networks. Adv Bioinformatics. 2011;2011:608295. doi: 10.1155/2011/608295. Epub 2011 Nov 29.

the number of input nodes, the Input(s), considered. The Rule(s) column indicate the fuzzy logic rule that associates the respective input node to the output node. The Error(E) column indicates the model's fit. Shown below is a sample output from the demo run:

```
> StartTime: Mon Jul 20 23:47:25 EDT 2020
> Search Parameters:
          inputFile = ./etc/projects/demo/inputs/exprsMat.txt
  maxNumberOfInputs = -1
     numberOfInputs = 3
   outputInRealtime = TRUE
            eCutOff = 0.6
useAllGenesAsOutput = FALSE
         iGeneStart = 1
           iGeneEnd = 14
        useParallel = TRUE
         outputFile = ./etc/projects/demo/outputs/runJFuzzy/exprsMat.1.14.3.TRUE.jfuz
     modelPhenotype = FALSE

Initiating...
Searching (Exhaustive Search)...
                 All Genes#: 14
   Output Nodes Considered#: 14
> Begin Search Result Table:
Output  NumberOfInput(s)    Input(s)    Rule(s) Error(E)
PTHLH   3   [KRT86, RUBCNL, CYS1]   [[1, 3, 3], [1, 3, 2], [2, 1, 1]]   0.6057812852061071
PTHLH   3   [LINC00707, RUBCNL, LINC00634]  [[1, 1, 1], [1, 2, 2], [2, 1, 1]]
    0.6395610849445412
PTHLH   3   [LINC00707, RUBCNL, LINC00634]  [[1, 1, 1], [1, 2, 3], [1, 1, 1]]
    0.6055562465480613
PTHLH   3   [LINC00707, RUBCNL, LINC00634]  [[1, 1, 1], [1, 3, 2], [2, 1, 1]]
    0.7444440833291
PTHLH   3   [LINC00707, RUBCNL, LINC00634]  [[1, 1, 1], [3, 2, 3], [2, 1, 1]]
    0.6035443648873104
PTHLH   3   [LINC00707, RUBCNL, LINC00634]  [[2, 1, 1], [1, 2, 2], [2, 1, 1]]
    0.664214821186895
PTHLH   3   [LINC00707, RUBCNL, LINC00634]  [[2, 1, 1], [1, 3, 2], [1, 1, 1]]
    0.6474161595899811
PTHLH   3   [LINC00707, RUBCNL, LINC00634]  [[2, 1, 1], [1, 3, 2], [2, 1, 1]]
    0.7321145800582836
  ...
```

```
C2orf78 3   [SERPINB7, CYS1, LINC00886] [[3, 3, 1], [1, 2, 3], [1, 3, 3]]
    0.6316025202379136
LINC00634   3   [LINC00886, GCNT4, MGAM]    [[1, 3, 3], [3, 2, 1], [2, 1, 1]]
    0.6000086433874239
LINC00634   3   [KRT86, LINC00886, GCNT4]   [[3, 1, 1], [2, 2, 3], [3, 1, 1]]
    0.6009588086189968
> End Search Result Table

...Done!
> Epilogue

   Started: 1595303245550: Mon Jul 20 23:47:25 EDT 2020
     Ended: 1595303288832: Mon Jul 20 23:48:08 EDT 2020
Total time: 43282 milliseconds; 0 min(s), 43 seconds.
```

#### The Utilities Module

jFuzzyMachine's Utilities Module consists of the 'Postprocessing' and the 'Add-ons' submodules. The postprocessing module consists of the 'Graph', 'Evaluation' and 'Dynamic Simulations' Units.

##### The Graph Unit

Outputs from the graph unit execution are placed in the runJFuzzUtils subdirectory. With regards to this demonstration, this would be the ./etc/projects/demo/outputs/runJFuzzy/runJFuzzUtils directory. These tab-delimited result files include:

- _demo_runJFuzzUtils.adj
- _demo_runJFuzzUtils.edg
- _demo_runJFuzzUtils.edg2
- _demo_runJFuzzUtils.fit
- _demo_runJFuzzUtils.fit2
- _demo_runJFuzzUtils.fre

The _demo_runJFuzzUtils.adj file is a directed graph adjacency matrix describing the connections in the inferred network. A connection between two nodes is indicated by 1 and 0 vice versa. Features in the rows are the inputs while those in columns are the output nodes. The _demo_runJFuzzUtils.edg and _demo_runJFuzzUtils.edg2 result files are about the same. Describing the edges in the inferred network, they both have the columns; From, To, Rule, and Weight in common. These correspond to the input node, output node, fuzzy logic rule associating the input with the output node, and estimated model fit respectively. The 'HashCode' column in the ".edg" file is only included for programmatic debugging. Likewise, the _demo_runJFuzzUtils.fit and _demo_runJFuzzUtils.fit2 result files are about the same. While the .fit2 reports all models above the specified fitCutOff in the jfuzzymachine.graph.config file, the .fit file reports only the best fitted model to each output node. Sampled outputs from the related demo run are shown below:

```
# _demo_runJFuzzUtils.fit
Output  NumberOfFittedModels    InputNodes(BestFit) Rules   Fit
C2orf78 7   [SERPINB7, CYS1, MGAM]  [[3, 3, 1], [1, 2, 3], [3, 1, 1]]   0.7246192458666514
LINC00634   2   [KRT86, LINC00886, GCNT4]   [[3, 1, 1], [2, 2, 3], [3, 1, 1]]   0.6009588086189968
    ...
```

```
# _demo_runJFuzzUtils.fit2
Output  InputNodes  Rules   Fits
C2orf78 [SERPINB7, CYS1, MGAM]  [[3, 3, 1], [1, 2, 3], [3, 1, 1]]   0.7246192458666514
C2orf78 [SERPINB7, CYS1, MGAM]  [[3, 2, 1], [1, 2, 3], [3, 1, 1]]   0.646429505593284
    ...
SERPINB7    [ROBO4, C2orf78, GCNT4] [[3, 2, 3], [3, 2, 1], [1, 3, 3]]   0.6002583965182628
SERPINB7    [ROBO4, C2orf78, LINC00634] [[1, 2, 3], [3, 1, 1], [2, 3, 1]]   0.6002285026737246
SERPINB7    [PTHLH, CYS1, C2orf78]  [[1, 3, 1], [3, 1, 1], [3, 2, 1]]   0.600143816707301
PTHLH   [LINC00707, RUBCNL, LINC00634]  [[1, 1, 1], [1, 3, 2], [2, 1, 1]]   0.7444440833291
PTHLH   [LINC00707, RUBCNL, LINC00634]  [[2, 1, 1], [1, 3, 2], [2, 1, 1]]   0.7321145800582836
    ...
```

The _demo_runJFuzzUtils.fre reports the frequency of the fuzzy rules evaluated in the inferred models with an estimated fit value above the fitCutOff. Please see Gormley et al [3] and Sokhansanj et al [4] for a detailed explanation of the rules.

[3]Gormley M, Akella VU, Quong JN, Quong AA. An integrated framework to model cellular phenotype as a component of biochemical networks. Adv Bioinformatics. 2011;2011:608295. doi: 10.1155/2011/608295. Epub 2011 Nov 29.

[4]Sokhansanj BA, Fitch JP, Quong JN, Quong AA. Linear fuzzy gene network models obtained from microarray data by exhaustive search. BMC Bioinformatics. 2004 Aug 10;5:108. doi: 10.1186/1471-2105-5-108.

##### The Evaluation Unit

- optparse

[5] R Core Team (2019). R: A language and environment for statistical computing. R Foundation for Statistical Computing, Vienna, Austria. URL https://www.R-project.org/.

[6] Huber W, Carey V, Gentleman R, et al. Orchestrating high-throughput genomic analysis with Bioconductor. Nat Methods 12, 115–121 (2015). https://doi.org/10.1038/nmeth.3252

[7] Gentleman RC, Carey VJ, Bates DM, Bolstad B, Dettling M, Dudoit S, Ellis B, Gautier L, Ge Y, Gentry J, Hornik K, et al. Bioconductor: open software development for computational biology and bioinformatics. Genome biology. 2004 Sep 1;5(10):R80.

- org.Hs.eg.db
- xtable
- igraph
- graph
- Rgraphviz
- pheatmap
- ReactomePA

To execute, simply run the following commands from within the jFuzzyMachine application working directory:

```
plugins/viz/rJFuzzyMachineUtilsExec.sh

plugins/viz/rJFuzzyMachineUtilsNetworkExec.sh
```

Example output figures, saved in the ./etc/projects/demo/outputs/plugins/viz/figs directory are presented below:

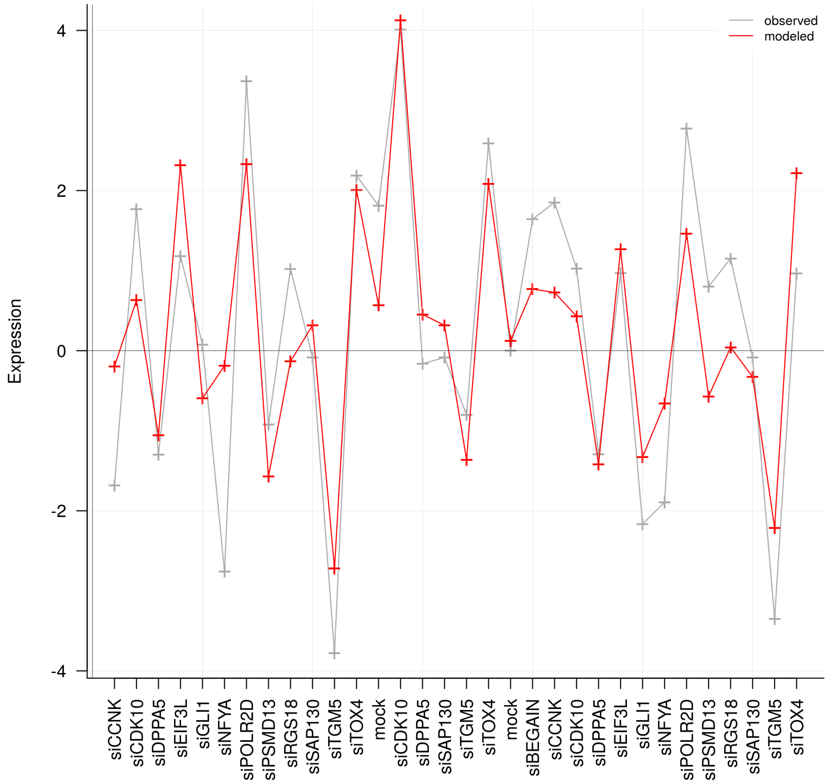

Figure 1: An Evaluation (Validation) Plot. A visual evaluation of predictions of a fitted model for a sample output node, the **C2orf78** gene. The estimated fit was 0.72. The input (regulatory) nodes were the genes *SERPINB7*, *CYS1*, and *MGAM*. The y-axis indicates the normalized expression values and the a-axis indicates the sample perturbations or treatment. Samples were reverse transfected vorinostat-resistant colon cancer, HCT-116, cell lines. Each sample was treated with the indicated small interferring ribonucleic acid (siRNA) to knockdown the respectively indicated gene products. The grey plot line shows the observed expression profile of the gene **C2orf78**, while the "red" line shows the predicted expression value from the expression of the regulators in the given data, and the rules associating the regulators to the output. The inferred patterns of regulation (rules) are indicated in Figure 3. It can apparently be appreciated that the fuzzy logic model is able to tease out trend in the dataset.

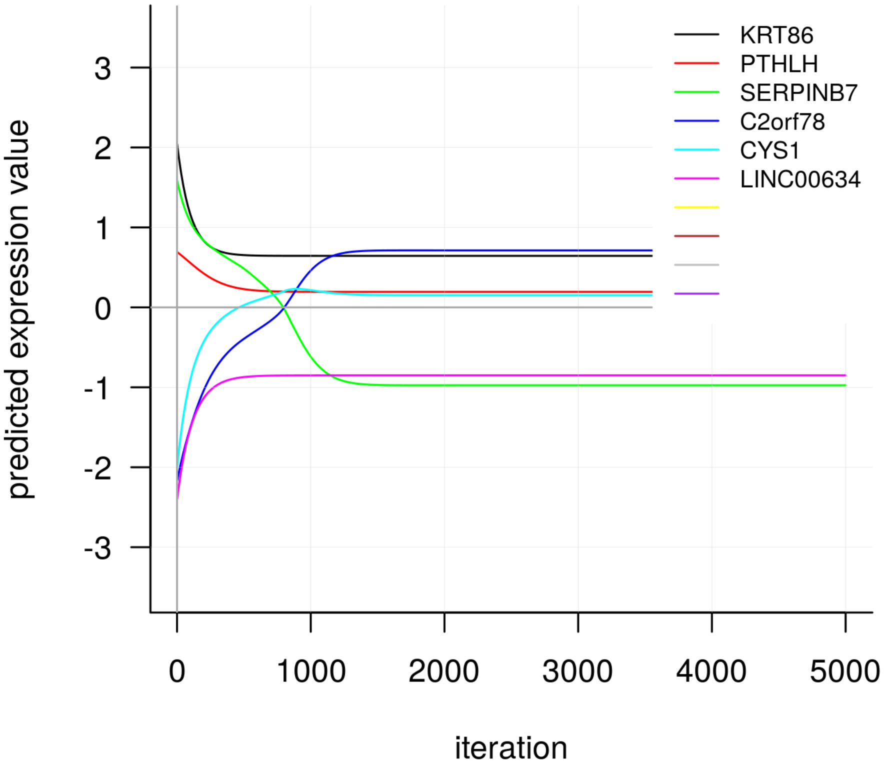

Figure 2: A Dynamic Simulation Plot. Having randomly chosen a sample (the 25th column sample) from the normalized expression matrix to provide initial values of expression, and given the best fitted models, the plot shows predicted expression values for the inferred outputs *KRT86*, *PTHLH*, *SERPINB7*, *C2orf78*, *CYS1* and *LINC00634* over 5000 iterations. It is appreciable that the inferred network achieves an equilibrium state at a little over 1000 iterations, when a change in predicted values tend to zero.

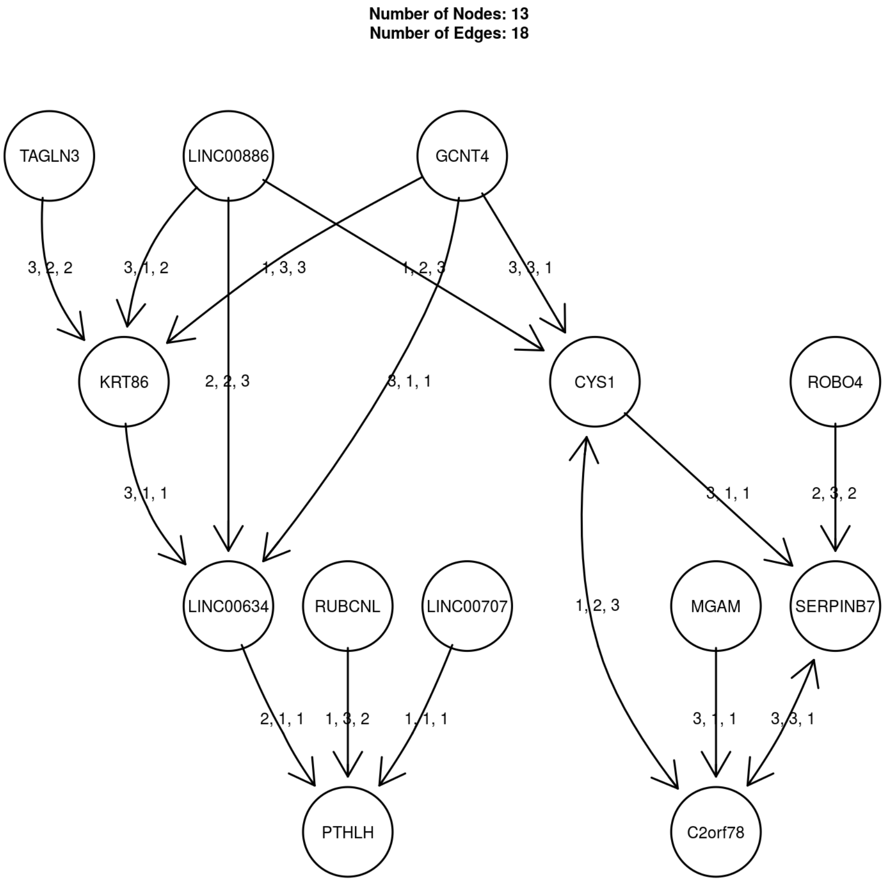

Figure 3: The Fuzzy Logic-based Regulatory Network Inferred. A composite regulatory network is inferred from the best fitted models for each node. The inferred network consists of 13 nodes (genes), and 18 edges (regulatory connections). The arrow heads indicate the regulatory direction from the input node to the output node. The edge labels, shown by the fuzzy rules, indicate the regulatory interaction. From Gormley et al, Rule configuration is the specification of if-then relationships between variables in fuzzy space. For example, an inhibitory relationship is represented by the rule vector $r = [r_1, r_2, r_3] = [3, 2, 1]$ (i.e., if input is low ($r_1$), then output is high (3); if input is medium ($r_2$), then output is medium (2), and if input is high ($r_3$), then ouput is low (1). From the composite regulatory network, the regulatory effect of the *MGAM* gene on the *C2orf78* gene is indicated by the rule $3, 1, 1$. This implies that when *MGAM* is low ($r_1$), *C2orf78* is high (3); when it is medium ($r_2$), *C2orf78* is low (1); and when *MGM* is high ($r_3$), *C2orf78* is low (1). Notice that the bi-directional relationship between the pair of genes *C2orf78–CYS1*, and *C2orf78–SERPINB7*
